# Use of Antibody Structural Information in Disease Prediction Models Reveals Antigen Specific B Cell Receptor Sequences in Bulk Repertoire Data

**DOI:** 10.1101/2024.12.10.627792

**Authors:** Onyekachi Nwogu, Kirandeep K. Gill, Carolina Moore, John W. Kroner, Wan-Chi Chang, Jeffrey Burkle, Mariana L. Stevens, Asel Baatyrbek kyzy, Emily R. Miraldi, Jocelyn M. Biagini, Ashley L. Devonshire, Leah Kottyan, Justin T. Schwartz, Amal H. Assa’ad, Lisa J. Martin, Sandra Andorf, Gurjit K. Khurana Hershey, Krishna M. Roskin

## Abstract

Convergent antibodies are highly similar antibodies elicited in multiple individuals in response to the same antigen. Convergent antibodies provide insight into shared immunological responses and show great promise as diagnostic biomarkers. They have typically been identified using methods that consider the amino acid sequence of the third complementarity-determining region (CDR3) of immunoglobulin heavy chain (IgH). In this study, we extend this definition of convergent antibodies to use structural information about the three IgH CDR regions (CDR1-3) and antibody isotype. We measure the performance of both definitions of convergence by their ability to predict disease status from bulk IgH sequencing data for two different disease cohorts: HIV infection and food allergen sensitization. We show that using predicted structural information outperforms prior approaches for the prediction of food sensitization status and performs on par for HIV infection status. Additionally, the use of structural information and antibody isotype enables better model interpretation and more immunological insights. The structurally convergent antibody groups driving HIV prediction accuracy are from known HIV-binding antibodies expressing the appropriate isotype. Thus, the incorporation of structural information can enable the identification of antigen specific antibody groups from bulk IgH sequencing data.

## Introduction

B cells develop and produce antibodies in response to antigens binding their B cell receptor (BCR). The BCR and the resulting antibody is composed of two polypeptide chains, known as the immunoglobulin (Ig) heavy-chain (IgH) and Ig light-chain, that are the result of rearrangement of variable (V), diversity (D), and junctional (J) gene segments during V(D)J recombination. This process produces a tremendous amount of combinatorial and junctional diversity in naive B cells. Each Ig chain contains six hypervariable loops called the complementarity determining regions (CDR 1-3). The heavy and light chains CDR1 and CDR2 loops are encoded by germline variable (V) genes segments, while the heavy-chain CDR3 stretches across the junctions of V(D)J recombination, making the heavy chain CDR3 the most diverse and “sufficient for most antibody specificities”[1]. When naive B cells interact with their antigen, this results in clonal expansion and somatic mutations that increases affinity[2]. Concurrently, B cells undergo class-switch recombination (CSR): genomic rearrangements that alter the constant region of the IgH chain, resulting in new isotypes enabling the elicitation of varying immune effector functions. CSR operates through the DNA excision, so while isotype switching can proceed sequentially along the IgH locus, it is irreversible. This highly diverse process of B cell development and maturation provides the ability to combat a vast array of pathogens and encodes current and past immune responses.

High-throughput sequencing technologies now enable the sequencing and monitoring of the antibody repertoire, including detection of convergent or shared antibodies: similar antibodies produced by multiple individuals when exposed to the same antigen. Convergent antibodies show great potential as diagnostic biomarkers of immune conditions [3, 4].

The traditional approach at identifying convergence is finding antibodies with highly similar amino acid sequences within IgH. This approach has been used to identify convergent antibodies for Dengue infection, H1N1 vaccination, HIV infection, tetanus immunization, lupus, COVID-19 and Type 1 Diabetes [3–7]. These methods use the germline identity of the IgH V- and J-gene segments and the amino acid sequence of the CDR3. We refer to this prior approach as the **VJ3 approach**.

Proteins with low amino acid sequence similarity can still have highly similar structures [8]. Therefore, only considering sequence similarity can miss antibodies that should be called convergent because of shared binding specificity, which is based on antibody structure. We present a new conceptualization of convergent antibodies that uses predicted antibody structural information. We call these **structurally convergent antibody groups (SCAGs)**.

Classification of antibodies based on stereotypic conformations, or *canonical classes*, has been discussed as early as the 1980’s for CDR1-2 [9, 10] and revised by several groups as recently as 2020 [11–15]. These methods have been expanded to include clustering of the heavy-chain CDR3s [16]. For IgH CDR1-2, we adopt the canonical class nomenclature from Wong et al. where the loops are grouped into one of 8 and 4 canonical classes respectively [17]. For the IgH CDR3, we used the clustering proposed by Choi et al. that annotates the CDR3 as belonging to one of 1169 classes [16]. Taking inspiration from recent works suggesting correlations between the variable and constant regions that affect antibody structure [19] and a dependence between expressed isotype and the V-, D-, and J-segment gene usage [18], we group sequences into the same SCAG if they have the same CDR1 and CDR2 canonical class, belong to the same CDR3 cluster, and express the same isotype.

We show the utility of the SCAGs definition of convergence by showing that in a disease prediction machine learning task, using SCAGs based features can greatly improve prediction and provide deeper immunological insights from bulk IgH sequencing data. We use two prediction tasks for this benchmarking: human immunodeficiency virus 1 (HIV-1) infection status and food allergen sensitization status. Additionally, we measure the effect on prediction accuracy of taking antibody isotype into account by building models that consider various isotype compartments and measuring their performance. We find that for the HIV prediction task, SCAG models, unlike VJ3 models, depend on SCAG features annotated with isotypes typically involved in the response to viral infections.

Our results show that prediction using antibody structural information greatly outperforms prior methods (for the food sensitization task) or was par with prior methods (for the HIV infection task) and the SCAG-based models prioritized features of the appropriate switched isotypes. For the HIV status prediction task, four of the top ten predictive SCAGs identified by the model were from known HIV-binding antibodies, including the highest weighted feature associated with the presence of HIV infection. Use of SCAGs allows the identification of antigen-specific antibody groups from bulk IgH sequencing data

## Results

### SCAG features outperform VJ3 features for the prediction of food sensitization status and performs on par for the prediction of HIV status

We use the presence or absence of convergent antibody groups as binary features in a logistic classification model. This additive linear model has been successfully used previously to predict HIV infection status[3]. We apply this framework to the SCAG and VJ3 definitions of convergent antibodies using subject-level stratified four-fold cross-validation for model training and testing, to prevent overfitting. We compare the performance of these features using receiver operating characteristic (ROC) curves and area under the ROC curve (AUC). We test the statistically significance of differences in AUC values using a two-sided DeLongs test[19]. While the VJ3 features used for the classification of HIV status recapitulated previously observed high performance[3] (Fig. 1A, red curve, AUC: 0.900), they perform poorly when used to statistically separate food sensitized subjects from those without food sensitization (Fig. 1B, red curve, AUC: 0.607). For the SCAGs models, the HIV classification model performed on par with the VJ3 model (Fig. 1A, green line, AUC: 0.889 vs. red line, AUC:0.900, DeLongs *p*=0.167), while the SCAGs food sensitization model significantly outperformed its VJ3 equivalent (Fig. 1B, green curve, AUC: 0.828 vs. red curve AUC: 0.607, DeLongs *p*=2.2×10^-16^).

**Figure 1:**
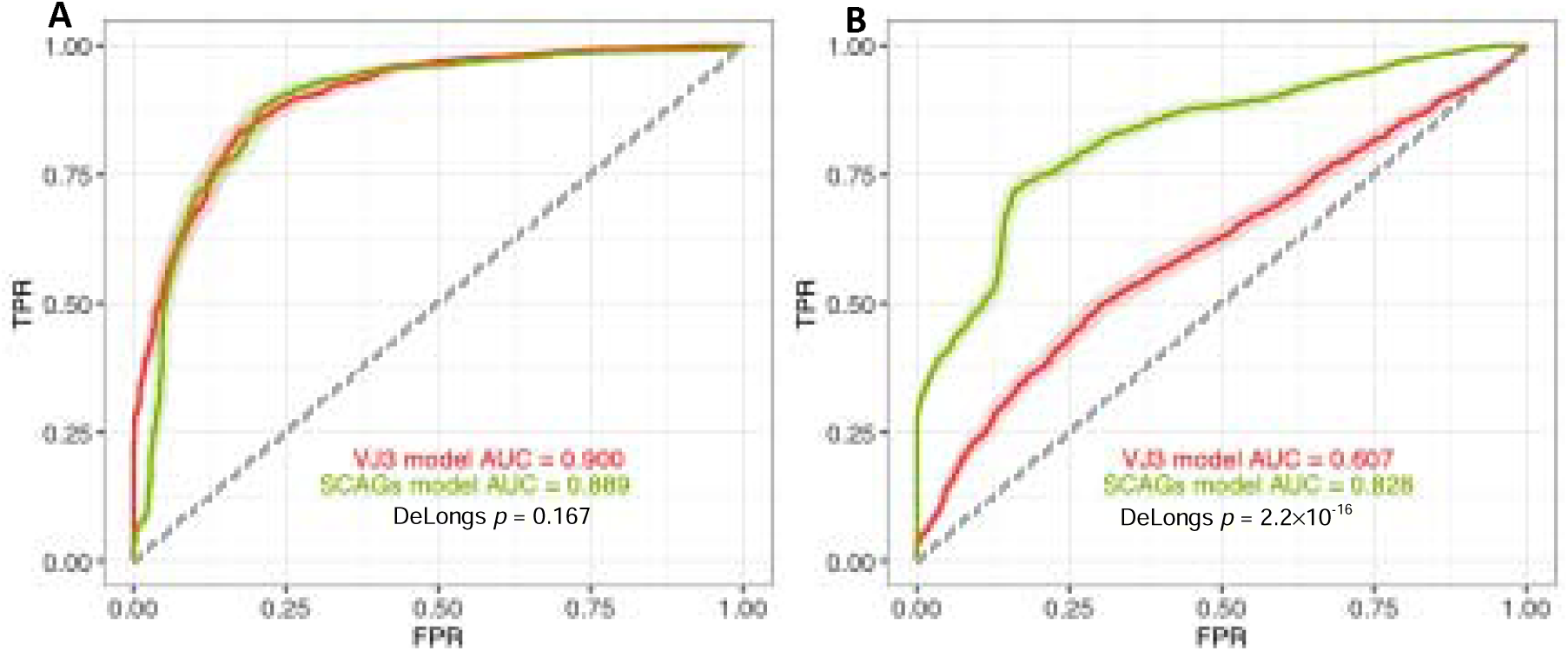
ROC curve and corresponding AUC values for VJ3 and SCAGs features. **(A)** HIV status prediction (HIV infected vs. HIV uninfected) test accuracy using VJ3 features (red curve) and using SCAGs features (green curve). **(B)** Food sensitization status prediction (food sensitized vs. not food sensitized) test accuracy using VJ3 features (red curve) and using SCAGs features (green curve). AUC scores are shown in red (VJ3) and green (SCAGs). True Positive Rate (TPR) and False Positive Rates (FPR) are calculated from the test fold of four-fold cross-validation and error ranges represent the standard error across all test folds. HIV status for participants in the HIV cohort was: 101 HIV infected and 43 not HIV infected. Sensitization status numbers for participants analyzed in the food sensitization cohort were: 70 sensitized and 83 non-sensitized.

### VJ3 and SCAG models focus on different isotype compartments, providing an explanation for task performance differences

To understand the large performance differences seen for the two different prediction tasks, we build variations of the VJ3 and SCAG models using additional forms of antibody isotype information.

Adding full isotype information to the VJ3 models, resulted in a non-significant change in accuracy on the HIV prediction task (Supplementary Fig. 1A, AUC 0.904 vs. 0.900, DeLongs *p*=0.641). Adding only the isotype-switched features (IgA, IgG, and IgE), i.e. excluding the naïve isotypes (IgM and IgD), resulted in a huge drop in accuracy on the HIV prediction task (Supplementary Fig. 1B, AUC 0.588 vs. 0.900, DeLongs *p*=2.2×10^-16^). On the other hand, removing the naïve isotypes from the SCAG model, had no significant effect on that model’s accuracy on the HIV prediction task (Supplementary Fig. 1C, AUC 0.891 vs. 0.889, DeLongs *p*=0.845). This suggests that the VJ3 models extract information primary from the naïve compartment, even in the absence of explicit isotype annotations, and this is sufficient for the high predictive accuracy on the HIV task. The SCAG models are able to achieve comparable accuracy on this task looking at distinct isotype compartments

On the food sensitization prediction task, adding isotype information to VJ3 model resulted in a significant but small improvement in accuracy (Supplementary Fig. 1D, AUC 0.641 vs. 0.607, DeLongs *p*=1.604×10^-8^). Adding only the isotype-switched features produced a small but non-significant increase in accuracy (Supplementary Fig. 1E, AUC 0.652 vs. 0.641, DeLongs *p*=0.216). For the SCAG model, removing naïve isotypes resulted in a significant but minor decrease in accuracy on the food sensitization task (Supplementary Fig. 1F, AUC 0.806 vs. 0.828, DeLongs *p*=4.609×10^-11^). This suggests that the majority of distinguishing antibody groups for the food sensitization prediction task are isotype-switched and thus harder for the VJ3 models to leverage, possibility explaining their poor performance.

### Model interpretation shows that SCAGs robustly predictive of HIV status use CDR3 conformations from known HIV-binding antibodies

For model interpretation, we extracted a set of SCAGs robustly predictive of HIV infection to understand the antibody groups the model was focusing on. As shown above, the naïve and isotype-switched compartments play a role in HIV status prediction, for VJ3 and SCAG respectively, thus we performed feature selection on data from all isotypes. Twenty-one HIV infection SCAGs were selected using random forest permutation importance followed by four-fold cross-validated LASSO regularization, a logical way to perform feature selection in the context of logistic regression [20], see Methods. Fourteen (67%) of the twenty-one SCAGs selected were positively associated with HIV infection (Supplementary Table S1) and all but one (93%) were of the IgG isotype. Four of the SCAGs associated with HIV infection shared CDR3 structural classification with known HIV-binding antibodies, including the highest weighted SCAG (Fig. 2, highlighted in red).

**Figure 2:**
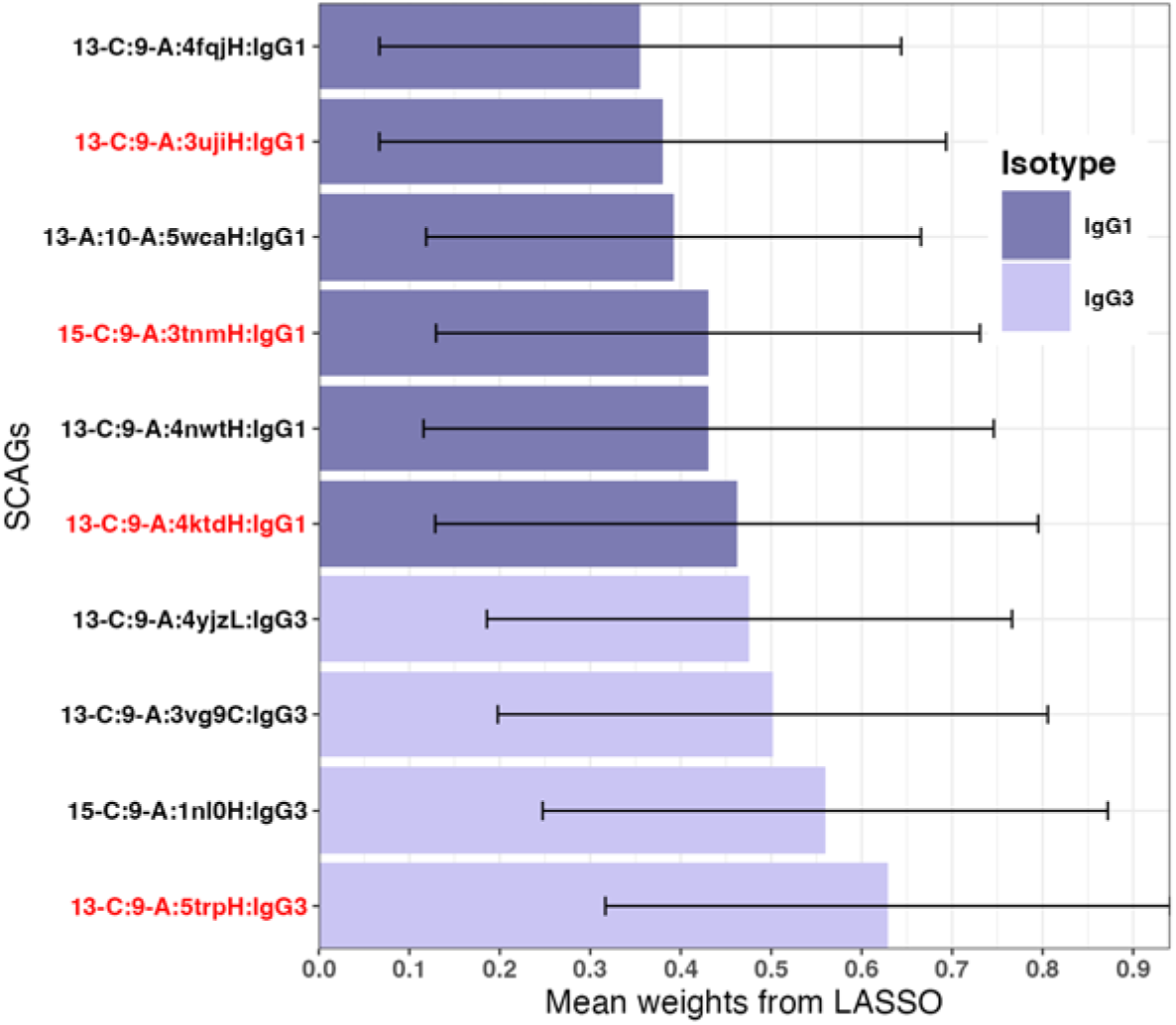
SCAGs regression weights for HIV status task after feature selection. Top ten SCAGs positively associated with HIV, out of twenty-one from four-fold cross-validated LASSO HIV prediction model (HIV infected vs. HIV uninfected). SCAGs that share a CDR3 structural cluster with known HIV-binding antibodies are highlighted in red. Features are labeled based on their assigned CDR1/2 canonical classes, CDR3 PDB chain, and expressed isotype. Error bars represent the range of weights (mean +/-standard deviation) across folds.

A similar model interpretation process found 9 SCAGs robustly predictive of food sensitization status (Supplementary Fig. 2 and Supplementary Table S2) with 7 being positively associated with food sensitization. Unlike for the HIV task, these SCAGs expressed a diverse set of isotypes.

### Model interpretation for food sensitization finds IgE isotype SCAG associated with non-sensitization

To interrogate the SCAG models when run on the food sensitization prediction task, we applied the same feature selection process described above. Since we found (see above) that the repertoire features discriminating food sensitized subjects from not food sensitized subjects reside in the isotype-switched compartment, we restricted feature selection to those expressing IgA, IgG, and IgE isotypes.

For our food sensitization cohort, seven robust class-switched features were selected. Five of which had positive weights and thus associated with food sensitization, with the remaining two having negative weights, associated with the absence of food sensitization (Supplementary Table S3). The second most predictive feature expressed IgE: *15-A:9-A:4qwwD:IgE,* a feature with a CDR1 belonging to canonical class 15-A, CDR2 belonging to class 9-A, and whose CDR3 is clustered with PDB chain 4qwwD, and is expressed as IgE (Fig. 3). Surprisingly, given the role IgE plays in atopic disease, this feature was associated with lack of food sensitization.

**Figure 3:**
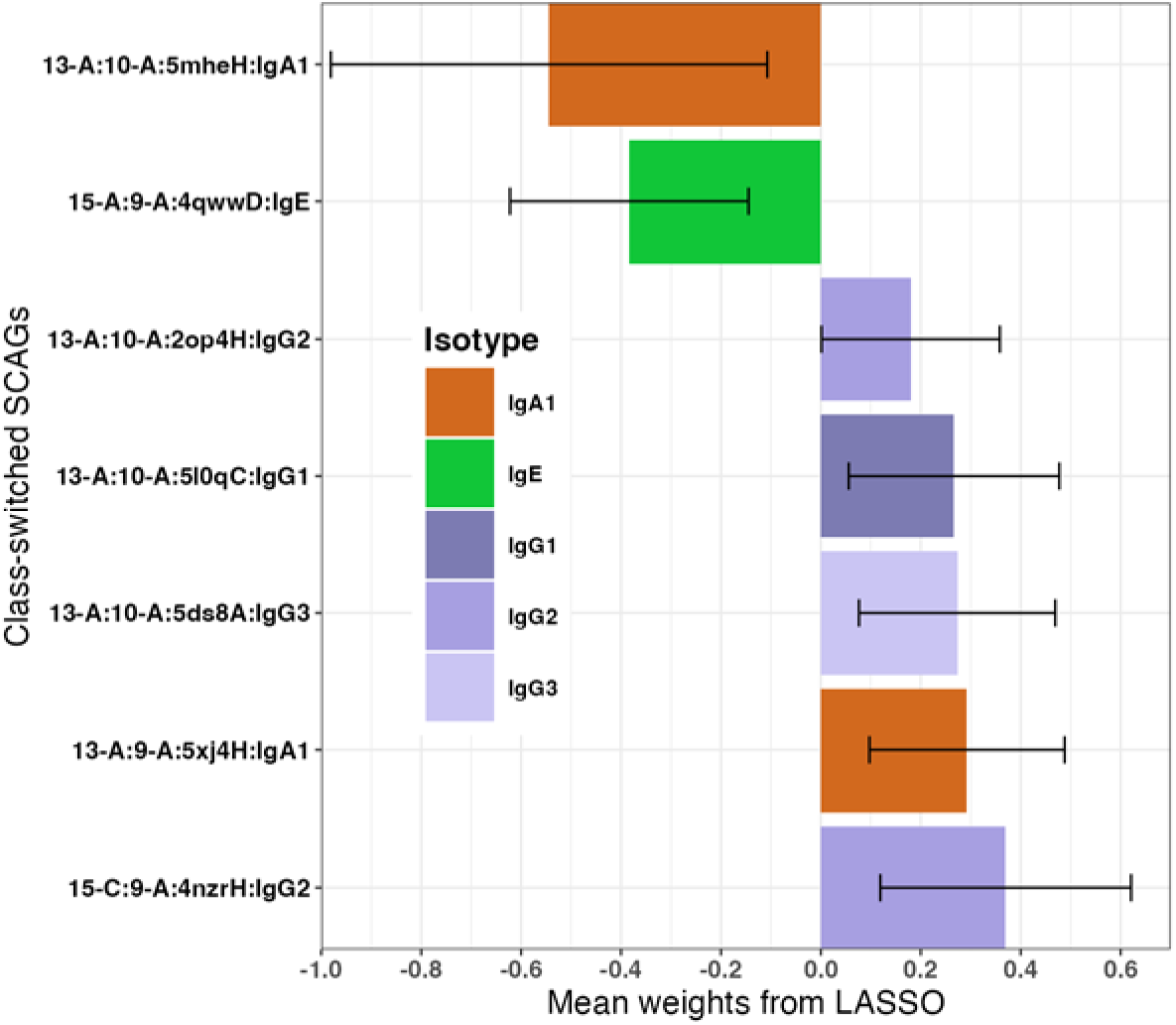
Class-switched SCAGs regression weights for food sensitization task after feature selection. Weights from four-fold cross-validated LASSO food sensitization status prediction model (food sensitized vs. non-food sensitized) using class-switched SCAGs features. The second most predictive group is of isotype class IgE and is associated with the absence of food sensitization. Features are labeled based on their assigned CDR1/2 canonical classes, CDR3 PDB chain, and expressed isotype. Error bars represent the range of weights (mean +/-standard deviation) across folds.

### IgE SCAG feature associated with lack of food sensitization is not associated with a specific allergen but does need to be of the IgE isotype

We investigated the association of SCAG feature *15-A:9-A:4qwwD:IgE* with non-sensitization to either peanut, egg, or milk (the most common food sensitizations in our dataset). A 1-sided Fisher’s Exact Test with correction for multiple hypothesis testing showed that *15-A:9-A:4qwwD:IgE* is associated with non-sensitization to both peanut (*adjusted p =1.56×10^-4^*) and egg (*adjusted p=1.55×10^-4^*), with a trend for milk being observed (*adjusted p=0.104*). Thus, the possible protective effects of 15-A:9-A:4qwwD:IgE are not restricted to one specific allergen.

We accessed if the “IgE-ness” of *15-A:9-A:4qwwD:IgE* is essential for its association with non-sensitization to food allergens with a 1-sided Fisher’s Exact Tests on all the related isotype forms of this SCAG. As expected, the IgE form is statistically significantly associated with non-sensitization to food allergens (*p=7.2×10^-7^*) but all the other related isotype-forms were not significantly associated with non-sensitization (*p>0.23*). We repeated this analysis for *15-A:9-A:4qwwD* (dropping the isotype restriction) and found no statistical association with non-food sensitization (*p>0.25*), further highlighting the importance of *15-A:9-A:4qwwD:IgE* being of the IgE isotype.

### SCAG features capture higher diversity in V-J segments and CDR3 sequences than VJ3 features

As expected, based on their definition, VJ3 features are very ridged in the IgH sequences they capture, this includes very limited complexity in the CDR3 amino acid sequences (Supplementary Table S4, Supplementary Fig. 3). Additional, VJ3 features are tied to the IGHV and IGHJ sequences and alleles captured in the ImMunoGeneTics (IMGT) nomenclature system [21]. Recent reports have found a large number of IGHV sequence alleles missing from the IMGT database [22]. By contrast, SCAG features capture a vast amount of diversity in IGHV and IGHJ segments used and particularly in the CDR3 amino acid sequence (Supplementary Fig. 4). SCAG features are also independent of the IMGT nomenclature system.

## Discussion

In this study, we investigated two different definitions of convergence antibody groups, VJ3 and SCAGs. VJ3 convergence groups are defined using the IMGT nomenclature system for V- and J-segments and sequence identity and have been used previously. Similar protein structures can include sequences that have low sequence similarity [23] and antibody structure is directly linked specificity. Structurally convergent antibody groups (SCAGs) are designed to leverage this fact and collect sequences with a tremendous amount of diversity (Supplementary Fig. 4). Additionally, based on new results suggesting a strong role played by antibody isotype (and the associated IGH constant region) on variable region selection and antibody specificity [24, 25], we include isotype as part of the SCAG definition. We benchmark these two definitions of convergent antibodies by measuring the prediction accuracy of machine learning models built using features derived from these definitions. We use two immunologically distinct prediction tasks: HIV status prediction and food sensitization status prediction.

We found that using VJ3 features to classify HIV infection using a logistic regression model, they perform excellently (Fig. 1A), replicating previous results [3]. Our novel SCAGs features performed on par in classifying HIV infection (Fig. 1A). On the food sensitization status prediction task, models built using SCAGs greatly outperformed models built with VJ3 features (Fig. 1B) achieving an AUC of 0.828.

In order to understand why SCAG features outperformed VJ3 features on the food sensitization task, we augmented VJ3 features with isotype information. This resulted only in a small accuracy increase (Supplementary Fig. 1D) suggesting that isotype is not the key difference between the VJ3 and SCAG models. Additionally, we performed ablation analysis by removing naïve isotype features from the models allowing us to inspect the effects of distinct isotype subsets on accuracy and correlate them to the immunology at play.

Models trained exclusively on class-switched features (i.e. IgA, IgG, or IgE) either performed on par with or experienced a minimal drop in accuracy relative to their full isotype counterparts. The exception was the VJ3 model on the HIV task, which showed a significant decrease in accuracy when the naïve isotypes (i.e. IgM and IgD) (Supplementary Fig. 1B) suggesting that the naïve compartment is critical to that model’s accuracy. HIV infection leads to immunodeficiency and abnormalities in lymphocyte populations[3, 26]. We hypothesize that this B cell deficiency manifests in the naïve compartment and is captured by VJ3 features. By contrast, both SCAGs feature models (class-switched only and full isotype models) perform similarly (Supplementary Fig. 1C) on the HIV prediction task. This suggests that SCAG models focus on the isotype-switched compartment, which would be expected to contain HIV-specific B cells.

To understand what SCAGs are driving prediction accuracy, we performed model interpretation by using cross-fold random forest feature importance filtering followed by LASSO logistic regression to obtain a reduced set of robust features for each classification task and convergent antibody approach (see Methods). On the HIV prediction task, this process yielded twenty-one HIV SCAGs, seventeen (81%) of which were of isotype class IgG1 (7 out of 21), IgG2 (4 out of 21) or IgG3 (6 out of 21), indicating that not only did our SCAGs model separate HIV infection status but it is focusing on the immune compartment (IgG) that produces antibodies against viruses (Fig. 3A, Supplementary Table S1). This is unlike the VJ3 with isotype model, where the naïve compartment was emphasized (Supplementary Table S5). Further evidence that SCAGs features are reflecting the underlying immunology is that, strikingly, 4 out of the 21 SCAGs belonged to the same CDR3 structural class as antibodies of known HIV antigen. Feature *13-C:9-A:5trpH*, the most predictive HIV SCAGs, shares its CDR3 structure with a known anti-HIV antibody binding the V1V2 and V3 loops on HIV-1 Env[27]. The ability of models using SCAGs features to identify known antigen-specific antibodies from bulk IgH repertoire data is an exciting development in immune repertoire analysis.

We also applied the same model interpretation pipeline to the isotype-switched SCAGs in food sensitization task. Interestingly, the second most predictive class-switched SCAG has the IgE isotype and is unexpectedly associated with absence of food sensitization (Fig. 3B, Supplementary Table S3). Only the IgE form of this SCAGs is associated with lack of food sensitization, with its non-isotype alternative not capturing this information. This observation would have been missed without the inclusion of isotype in our convergence definitions. We hypothesis that the IgE antibodies belonging to these features might be protective against food sensitization. A study by Keswani et al. identified a peanut specific IgG4 antibody capable of blocking anaphylaxis in mice [28]. The putatively protective antibody structure we have identified seems to be required to be IgE for its protective effects. Additionally, it seems to be protective for 2 out of the 3 common food allergens in our cohort. Taken together, this suggests that the protective interactions involve IgE-FcεR1 interactions rather than IgE-allergen interactions. These findings further demonstrate the immunological knowledge that adding isotypes to our convergence antibody definitions provides.

Although most allergies are lifelong, IgE producing plasma B cells are short lived[29], while IgE memory B cells are rare[30, 31]. Several studies have hypothesized that IgE memory to allergen specific antigens is stored as IgG memory B cells[32, 33]. This may explain the presence of majority of IgG SCAGs (57%) associated with food sensitization (Fig. 3B). These IgG B cells may be poised to class switch to IgE and become short-lived plasma blasts.

While our clinical cohorts encompass broad biological conditions, SCAGs provide a computational framework capable of filtering complex repertoires down to testable immunological hypotheses that serve as building blocks for uncovering vulnerable binding sites. For instance, we observe a potentially protective IgE antibody, whose protection seems to be dependent on being of the IgE isotype. We also identify predominantly IgG features associated with food sensitization, in line with recent hypotheses that allergic specific IgG B cells may hold memory for IgE antibodies. While SCAGs did not markedly improve performance for the prediction of HIV infection status, incorporating that information into the model did recapitulate know immunology. From completely untargeted sequencing of the IgH repertoire from the periphery, we identified four predictive HIV SCAG features that share the same CDR3 structure as known HIV-binding antibodies, with the majority belonging to IgG subclasses (predominantly IgG1 and IgG3), in line with immunoglobulin response to HIV and viral infections. Together, these findings highlight the tremendous additional immunological information gained by incorporating structural and isotype information into interpretable machine learning methods for the prediction of diseases from bulk IgH sequencing data, successfully narrowing millions of candidate sequences down to public lineages suitable for targeted experimental validation.

## STAR Methods

### Key resources table

| REAGENT or RESOURCE | SOURCE | IDENTIFIER |
|---|---|---|
| **Biological samples** |  |  |
| AIRR-seq data from atopic study | NIH NLM BioProject | PRJNA857098 |
| AIRR-seq data from HIV study | NIH NLM BioProject | PRJNA486667 |
| **Chemicals, peptides, and recombinant proteins** |  |  |
| Histopaque-1077 | Sigma-Aldrich |  |
| Random Hexamers | Invitrogen |  |
| Agarose (for gel extraction) | Standard Commercial Supplier |  |
| **Critical commercial assays** |  |  |
| SepMate column system kit | StemCell Technologies |  |
| AllPrep column purification kit | Qiagen |  |
| SuperScript III First-Strand Synthesis System | Invitrogen |  |
| Platinum PCR Master Mix kit | Applied Biosystems |  |
| Qiagen Multiplex PCR kit | Qiagen |  |
| QIAquick Gel Extraction kit | Qiagen |  |
| MiSeq Reagent Kit (600-cycle) | Illumina |  |
| **Oligonucleotides** |  |  |
| BIOMED-2 IGHV framework primers with P5/P7 adapters | BIOMED-2 / Custom Synthesis | Ref [34, 35] |
| Isotype-specific CH1 primers with P5/P7 adapters | Custom Synthesis | Ref [34, 35] |
| **Software and algorithms** |  |  |
| IgBLAST | Ye et al., 2013 | https://www.ncbi.nlm.nih.gov/igblast/ |
| FLASH | Magoc et al., 2011 | https://www.psc.edu/resources/software/flash/ |
| MMseqs2 | Steinegger et al., 2017 | https://github.com/soedinglab/MMseqs2 |
| SAAB+ | Kovaltsuk et al., 2020 | https://github.com/oxpig/saab_plus |
| SCALOP | Wong et al.,2018 | https://github.com/oxpig/SCALOP |
| FREAD | Choi et al., 2010 | https://github.com/oxpig/saab_plus |
| ggplot2, version 3.5.1 | Wickham et al., 2016 | https://cran.r-project.org/web/packages/ggplot2/index.html |
| ggseqlogo, version 0.2 | Wagih et al., 2017 | https://github.com/omarwagih/ggseqlogo |
| pROC version, 1.18.5 | Robin et al., 2011 | https://xrobin.github.io/pROC/ |
| scikit-learn version, 1.0.2 | Pedregosa et al., 2011 | https://github.com/scikit-learn/scikit-learn |
| **Other** |  |  |
| Illumina MiSeq | Illumina | Ref [36] |
| Next-Generation Sequencing Core | CCHMC DNA Core |  |

### HIV cohort study design

Participants were recruited in a multi-site study of HIV-infection that included an uninfected control arm as previously described in [3, 26]. In brief, virus controllers and subjects who did not give informed consent were excluded from the study and infected and uninfected subjects were match using propensity score-matching with age, sex, and country of origin as variables. Subjects with HIV infection and uninfected controls were predominantly African[26]. The characteristics of this cohort are shown in Supplementary Table S6.

### Atopic cohort study design

The food sensitization dataset consisted of high throughput sequencing of rearranged IgH genes from peripheral B cells of 153 children with atopic dermatitis (AD) participating in the Mechanism of Progression of Atopic Dermatitis to Asthma in children (MPAACH) study [37, 38]. Inclusion and exclusion criteria for MPAACH are described in [38] but in brief subjects were born at gestation of ≥36 weeks, aged 1–2 years on enrollment, had AD, and no co-morbid lung condition or dependence on immunosuppression or oral steroids. All subjects or their legal guardian gave informed consent prior to participation. Allergic sensitization was measured on the day of blood draw by skin prick testing (SPT) on a panel of all 13 foods and 11 aeroallergens[38]. The characteristics of the cohort are shown in supplementary Table S7.

### IgH sequencing libraries

Sequencing libraries were generated as described in [3] and [39]. In short peripheral blood mononuclear cells (PBMCs) were isolated from whole blood and complementary DNA (cDNA) was generated from total RNA using random hexamers, followed by PCR amplification of IGH rearrangements using multiplex primers targeting IGHV gene segments using the BIOMED-2 FR1 primers and isotype-specific primers located in CH1[34, 35, 40]. Each isotype was amplified separately to decrease cross-priming and the resulting libraries were pooled and purified by agarose gel electrophoresis and gel extracted before sequencing on the Illumina MiSeq.

### IgH sequence annotation

High-throughput data was processed as previously described[3, 40]. In summary, 300bp paired- end reads were merged using FLASH[41] and reads where mapped to samples using barcodes. The V-,D-, and -J segments, framework and CDR3 regions were identified using IgBLAST(version 1.16.0)[42]. Germline sequences for the V-,D-, and-J segments were obtained from the IMGT database (version 202232-2)[21]. Each sequence underwent quality filtering whereby only productive reads having a CDR3 region, V segment alignment score of at least 70 and J segment alignment score of at least 26. Each transcript’s isotype was determined by matching it to a database of constant region gene sequences upstream from the primer[21].

### Clonal inferences

Clonal relationships were assigned as previously described, using MMseqs2 (Sep. 2019 build) [43] to cluster the sequences into clones having the same V- and J-segments (without considering the allele), equal CDR-H3 length, and at least 90% CDR-H3 nucleotide identity [35].

### VJ3 convergent groups features

VJ3 convergent antibodies were identified as follows. CDR3 sequences for each cohort (HIV and Atopy) were clustered using MMseqs2 (Sep. 2019 build)[43], whereby each cluster contained CDR3 sequences having equal lengths, annotated with the same V-and J-segments (excluding alleles), 80% bidirectional sequence coverage, and 90% CDR3 amino acid sequence identity. For each cluster, consensus amino acid sequence of the CDR3 was calculated. A subject is said to have antibody sequences belonging to a convergent group if they express sequences annotated with the same V- and J-segments as the convergent group and a 90% CDR3 amino acid sequence identity match to the consensus CDR3 sequence of the group.

### Structurally convergent antibody group features

IgH sequences were processed with SAAB+[16], an analysis pipeline which annotates BCR amino acid sequences with structural information using SCALOP[17], designating canonical classes of CDR1 and CDR2 regions and FREAD[16] to assign CDR3 sequences to predicted structural clusters.

SCAG convergent antibodies were defined as follows: IgH sequences having the same CDR1, CDR2 canonical classes, CDR3 loop structures as annotated by SAAB+ and expressed isotype.

### Predictive model using VJ3 and SGAGs features

Binary feature matrices of identified convergent features from all the methods explored were created, with columns representing the convergent group and rows representing subjects. Elements of the matrices are 1 if the subject expresses a sequence that is a member of a convergent group and 0 otherwise. Four-fold stratified cross-validation at the subject level was employed to avoid overfitting during model fitting and testing when assessing the performance of our models. Logistic and LASSO (described below) regression models using the scikit-learn library[44] in python was applied in building models to predict HIV and food sensitization status. For the logistic regression models, only features in the train subsets shared in at least 11 subjects are kept.

### Model interpretation

A two-phase approach was utilized for to select features for model interpretation and exploration. First features were selected if their random forest permutation importance value was above the absolute value of the most negative feature importance[45]. Features with negative importance, i.e. features that *improve* model performance when randomly permutated, represent the threshold for random importance[46]. The remaining features were used in the second phase, where several cross-validated (repeated four-fold cross-validation) LASSO models were fitted with varying regularization strengths using the scikit-learn package (version 1.0.2). Robust features were selected as those whose mean weight plus or minus their standard deviation across all folds did not intersect zero. A regularization strength of the LASSO model was selected based on the local maxima in the number of robust features (Supplementary Fig. 5).

## Statistical analysis

Statistical analysis and graphs were carried out using R statistical language (version 4.4.0), in R studio (2024.04 release) and python (version 3.4.10). The logistic regression analysis was done using the scikit-learn package (version 1.0.2) and the LogisticRegression function in python. LASSO regression was applied using the LogisticRegression function with ‘l1’ penalty. Area under the curve (AUC) was calculated using the pROC package (version 1.18.5) in R[47]. Fisher’s Exact and DeLong Tests were implemented using the fisher.test function and pROC’s roc.test function in R respectively. A P-value of 0.05 was considered significant.

## Visualizations

Bar, tile and CDR3 sequence logo plots were created using the ggplot2 (version 3.5.1) package[48] and the ggseqlogo (version 0.2) package[49] in R. Receiver operating characteristics curves (ROC) were generated using the ggroc function in using the pROC (version 1.18.5) package[47] in R.

## Acknowledgments

The authors thank all the children and their families who participated in the MPAACH cohort, the Schubert Research Clinic of Cincinnati Children’s Hospital Medical Center (CCHMC) for assistance with research participants, the CCHMC Integrated Genomics and Microbiome Sequencing Facility (RRID:SCR_022630) and in particular Brian Quinn for help optimizing library construction), the CCHMC Information Services for Research (IS4R, RRID:SCR_022622) group for hosting the data storage and processing infrastructure.

## Data and materials availability

All food sensitization data associated with this study are available from the NIH Short Read Archive (SRA) under BioProject <u>PRJNA857098</u>. Data from the HIV cohort can be found in the BioProject <u>PRJNA486667</u>. All python and R code is available upon request.

## References

1. Xu, J.L. and M.M. Davis, Diversity in the CDR3 region of V(H) is sufficient for most antibody specificities. Immunity, 2000. 13(1): p. 37–45.

2. Frost, S.D., et al., Assigning and visualizing germline genes in antibody repertoires. Philos Trans R Soc Lond B Biol Sci, 2015. 370(1676).

3. Roskin, K.M., et al., Aberrant B cell repertoire selection associated with HIV neutralizing antibody breadth. Nat Immunol, 2020. 21(2): p. 199–209.

4. Zaslavsky, M.E., et al., Disease diagnostics using machine learning of B cell and T cell receptor sequences. Science, 2025. 387(6736): p. eadp2407.

5. Parameswaran, P., et al., Convergent antibody signatures in human dengue. Cell Host Microbe, 2013. 13(6): p. 691–700.

6. Jackson, K.J., et al., Human responses to influenza vaccination show seroconversion signatures and convergent antibody rearrangements. Cell Host Microbe, 2014. 16(1): p. 105–14.

7. Burckert, J.P., et al., Functionally Convergent B Cell Receptor Sequences in Transgenic Rats Expressing a Human B Cell Repertoire in Response to Tetanus Toxoid and Measles Antigens. Front Immunol, 2017. 8: p. 1834.

8. Brenner, S.E., et al., Understanding protein structure: using scop for fold interpretation. Methods Enzymol, 1996. 266: p. 635–43.

9. Chothia, C. and A.M. Lesk, Canonical structures for the hypervariable regions of immunoglobulins. J Mol Biol, 1987. 196(4): p. 901–17.

10. Chothia, C., et al., Conformations of immunoglobulin hypervariable regions. Nature, 1989. 342(6252): p. 877–83.

11. Kelow, S.P., J. Adolf-Bryfogle, and R.L. Dunbrack, Hiding in plain sight: structure and sequence analysis reveals the importance of the antibody DE loop for antibody-antigen binding. MAbs, 2020. 12(1): p. 1840005.

12. Adolf-Bryfogle, J., et al., PyIgClassify: a database of antibody CDR structural classifications. Nucleic Acids Res, 2015. 43(Database issue): p. D432–8.

13. Dunbar, J., et al., SAbDab: the structural antibody database. Nucleic Acids Res, 2014. 42(Database issue): p. D1140–6.

14. North, B., A. Lehmann, and R.L. Dunbrack, Jr., A new clustering of antibody CDR loop conformations. J Mol Biol, 2011. 406(2): p. 228–56.

15. Nowak, J., et al., Length-independent structural similarities enrich the antibody CDR canonical class model. MAbs, 2016. 8(4): p. 751–60.

16. Choi, Y. and C.M. Deane, FREAD revisited: Accurate loop structure prediction using a database search algorithm. Proteins, 2010. 78(6): p. 1431–40.

17. Wong, W.K., et al., SCALOP: sequence-based antibody canonical loop structure annotation. Bioinformatics, 2019. 35(10): p. 1774–1776.

18. Montamat-Garcia, G., et al., Human primary antibody response to vaccination follows a partially sequential class-switching program with a checkpoint at IGHG2. Cell Rep Med, 2026. 7(7): p. 102848.

19. DeLong, E.R., D.M. DeLong, and D.L. Clarke-Pearson, Comparing the areas under two or more correlated receiver operating characteristic curves: a nonparametric approach. Biometrics, 1988. 44(3): p. 837–45.

20. Tibshirani, R., Regression shrinkage and selection via the Lasso. Journal of the Royal Statistical Society Series B-Statistical Methodology, 1996. 58(1): p. 267–288.

21. Lefranc, M.P., et al., IMGT(R), the international ImMunoGeneTics information system(R) 25 years on. Nucleic Acids Res, 2015. 43(Database issue): p. D413–22.

22. Jana, U., et al., The human IG heavy chain constant gene locus is enriched for large structural variants and coding polymorphisms that vary among human populations. Cell Genom, 2026. 6(1): p. 101058.

23. Holm, L. and C. Sander, Mapping the protein universe. Science, 1996. 273(5275): p. 595–603.

24. McConnell, S.A. and A. Casadevall, New insights into antibody structure with implications for specificity, variable region restriction and isotype choice. Nat Rev Immunol, 2025.

25. McConnell, S.A. and A. Casadevall, Immunoglobulin constant regions provide stabilization to the paratope and enforce epitope specificity. J Biol Chem, 2024. 300(6): p. 107397.

26. Moody, M.A., et al., Immune perturbations in HIV-1-infected individuals who make broadly neutralizing antibodies. Sci Immunol, 2016. 1(1): p. aag0851.

27. Bonsignori, M., et al., Staged induction of HIV-1 glycan-dependent broadly neutralizing antibodies. Sci Transl Med, 2017. 9(381).

28. Keswani, T., et al., Neutralizing IgG(4) antibodies are a biomarker of sustained efficacy after peanut oral immunotherapy. J Allergy Clin Immunol, 2024.

29. Yang, Z., B.M. Sullivan, and C.D. Allen, Fluorescent in vivo detection reveals that IgE(+) B cells are restrained by an intrinsic cell fate predisposition. Immunity, 2012. 36(5): p. 857–72.

30. He, J.S., et al., Biology of IgE production: IgE cell differentiation and the memory of IgE responses. Curr Top Microbiol Immunol, 2015. 388: p. 1–19.

31. He, J.S., et al., The distinctive germinal center phase of IgE+ B lymphocytes limits their contribution to the classical memory response. J Exp Med, 2013. 210(12): p. 2755–71.

32. Hoof, I., et al., Allergen-specific IgG(+) memory B cells are temporally linked to IgE memory responses. J Allergy Clin Immunol, 2020. 146(1): p. 180–191.

33. Ota, M., et al., The memory of pathogenic IgE is contained within CD23 (+) IgG1 (+) memory B cells poised to switch to IgE in food allergy. bioRxiv, 2023.

34. van Dongen, J.J., et al., Design and standardization of PCR primers and protocols for detection of clonal immunoglobulin and T-cell receptor gene recombinations in suspect lymphoproliferations: report of the BIOMED-2 Concerted Action BMH4-CT98-3936. Leukemia, 2003. 17(12): p. 2257– 317.

35. Roskin, K.M., et al., IgH sequences in common variable immune deficiency reveal altered B cell development and selection. Sci Transl Med, 2015. 7(302): p. 302ra135.

36. Gill, K., et al., B cell repertoire in children with skin barrier dysfunction supports altered IgE maturation associated with allergic food sensitization. bioRxiv, 2023.

37. Gill, K., et al., B cell repertoire in children with skin barrier dysfunction supports altered IgE maturation associated with allergic food sensitization. bioRxiv, 2023.

38. Biagini Myers, J.M., et al., Events in Normal Skin Promote Early-Life Atopic Dermatitis-The MPAACH Cohort. J Allergy Clin Immunol Pract, 2020. 8(7): p. 2285–2293 e6.

39. Gill, K.K., et al., B-Cell Repertoire of Children With Atopic Dermatitis Exhibits Altered IgE Maturation Associated With Allergic Food Sensitization. Allergy, 2026. 81(2): p. 513–523.

40. Nielsen, S.C.A., et al., Shaping of infant B cell receptor repertoires by environmental factors and infectious disease. Sci Transl Med, 2019. 11(481).

41. Magoc, T. and S.L. Salzberg, FLASH: fast length adjustment of short reads to improve genome assemblies. Bioinformatics, 2011. 27(21): p. 2957–63.

42. Ye, J., et al., IgBLAST: an immunoglobulin variable domain sequence analysis tool. Nucleic Acids Res, 2013. 41(Web Server issue): p. W34–40.

43. Steinegger, M. and J. Soding, MMseqs2 enables sensitive protein sequence searching for the analysis of massive data sets. Nat Biotechnol, 2017. 35(11): p. 1026–1028.

44. Fabian Pedregosa, G.V., Alexandre Gramfort, Vincent Michel, Bertrand Thirion, Olivier Grisel, Mathieu Blondel, Peter Prettenhofer, Ron Weiss, Vincent Dubourg, Jake Vanderplas, Alexandre Passos, David Cournapeau, Matthieu Brucher, Matthieu Perrot, Édouard Duchesnay, Scikit-learn: Machine Learning in Python. Journal of Machine Learning Research, 2011. 12: p. 2825––2830.

45. Wiener, A.L.a.M., Classification and Regression by randomForest. R News, 2002. 2(3): p. 18–22.

46. Chinthrajah, R.S., et al., Development of a tool predicting severity of allergic reaction during peanut challenge. Ann Allergy Asthma Immunol, 2018. 121(1): p. 69–76 e2.

47. Robin, X., et al., pROC: an open-source package for R and S+ to analyze and compare ROC curves. BMC Bioinformatics, 2011. 12: p. 77.

48. Wickham, H., ggplot2: Elegant Graphics for Data Analysis. 2016: Springer-Verlag New York.

49. Wagih, O., ggseqlogo: a versatile R package for drawing sequence logos. Bioinformatics, 2017. 33(22): p. 3645–3647.

