## Supplemental Figures and Tables for "Use of Antibody Structural Information in Disease Prediction Models Reveals Antigen Specific B Cell Receptor Sequences in Bulk Repertoire Data"

**Supplementary Table S1: Characteristics of stable SCAGs features selected by cross-validated LASSO model for the prediction of HIV infection status.** HIV-Pos and HIV-Neg signify the number of infected and uninfected subjects expressing sequences belonging to any of the 21 SCAGs features. Total represents the total number of subjects in our study cohort expressing sequences belonging to each of the 21 SCAGs features. For each stable SCAGs feature, their mean weights and standard deviation across folds was calculated. A description of the antibody source of the CDR3 structure is also shown.

| **SCAGs Features** | **HIV-Pos** | **HIV-Neg** | **Total** | **Weights mean** | **Std. dev of weights** | **CDR3 Source Description** |
| --- | --- | --- | --- | --- | --- | --- |
| 13-C:9-A:5trpH:IgG3 | 38 | 3 | 41 | 0.628 | 0.312 | DH272, lineage member of DH270 HIV bNAb |
| 15-C:9-A:1nl0H:IgG3 | 39 | 2 | 41 | 0.559 | 0.312 | Conformation-specific anti-factor IX antibody |
| 13-C:9-A:3vg9C:IgG3 | 73 | 10 | 83 | 0.501 | 0.303 | Monoclonal antibody against human A(2A) AR |
| 13-C:9-A:4yjzL:IgG3 | 74 | 13 | 87 | 0.4759 | 0.290 | Human H2526 in complex with influenza hemagglutinin H1 |
| 13-C:9-A:4ktdH:IgG1 | 91 | 22 | 113 | 0.461 | 0.333 | Non-human primate HIV vaccine-elicited GE136 |
| 13-C:9-A:4nwtH:IgG1 | 62 | 9 | 71 | 0.4309 | 0.315 | anti-human β-nerve growth factor antibody |
| 15-C:9-A:3tnmH:IgG1 | 87 | 20 | 107 | 0.430 | 0.300 | A32 Fab, ADCC mediating anti-HIV-1 antibody |
| 13-A:10-A:5wcaH:IgG1 | 70 | 10 | 80 | 0.392 | 0.273 | VRC 315 27-1C08, Influenza bNAb |
| 13-C:9-A:3ujiH:IgG1 | 87 | 17 | 104 | 0.380 | 0.313 | Anti-HIV-1 V3 Fab 2558 |
| 13-C:9-A:4fqjH:IgG1 | 76 | 12 | 88 | 0.355 | 0.288 | CR8071, influenza B hemagglutinin antibody |
| 13-C:9-A:1nl0H:IgG3 | 46 | 5 | 51 | 0.339 | 0.300 | Conformation-specific anti-factor IX antibody |
| 13-A:10-B:3ln9A:IgG1 | 77 | 15 | 92 | 0.279 | 0.260 | Fibril-specific B10 antibody |
| 15-C:9-A:2yc1D:IgD | 80 | 21 | 101 | 0.234 | 0.223 | 9004G, Cn2 toxin binding antibody |
| 13-A:10-A:4krmL:IgG3 | 31 | 0 | 31 | 0.185 | 0.184 | Inhibitory EGFR nanobody/VHH domain |
| 13-A:9-A:1bgxH:IgA1 | 23 | 21 | 44 | -0.355 | 0.247 | TP7, TAQ polymerase binding antibody |
| 13-A:10-A:6be3H:IgG2 | 47 | 31 | 78 | -0.400 | 0.286 | F598, GlcNAc binding antibody |
| 13-A:10-B:2aepH:IgG2 | 32 | 27 | 59 | -0.479 | 0.285 | antibody binding influenza neuraminidase |
| 13-A:10-A:6cmiD:IgM | 55 | 32 | 87 | -0.512 | 0.252 | anti-Hendra virus attachment glycoprotein G antibody |
| 13-A:10-B:1a6vJ:IgG2 | 23 | 22 | 45 | -0.544 | 0.272 | Fv complexed with acetate compound |
| 13-A:10-A:4qhuB:IgG2 | 26 | 26 | 52 | -0.676 | 0.292 | Fab6785, IL-17A binding antibody |
| 13-A:10-B:2yc1D:IgA1 | 59 | 36 | 95 | -0.708 | 0.312 | 9004G, Cn2 toxin binding antibody |

**Supplementary Table S2: Characteristics of stable SCAGs features selected by cross-validated LASSO model for the prediction of food sensitization status.** Sensit and Non-Sensit signify the number of food sensitized subjects and subjects without a food sensitization expressing sequences belonging to any of the 9 SCAGs features. Total represents the total number of subjects in our study cohort expressing sequences belonging to each of the 9 SCAGs features. For each stable SCAGs feature, their mean weights and standard deviation across folds were calculated. A description of the antibody source of the CDR3 structure is also shown.

| SCAGs Features | Sensit | Non-Sensit | Total | Coefficient mean | Std. dev of weights | CDR3 Source Description |
| --- | --- | --- | --- | --- | --- | --- |
| 15-C:9-A:4nzrH:IgG2 | 41 | 10 | 51 | 0.445 | 0.281 | PGT135, HIV binding antibody (structure not bound to antibody) |
| 13-A:9-A:5xj4H:IgA1 | 50 | 23 | 73 | 0.424 | 0.217 | Complex structure of durvalumab-scFv/PD-L1 |
| 13-A:10-B:4m43H:IgD | 47 | 20 | 67 | 0.385 | 0.241 | Fab 523-11, anti-rabies antibody |
| 13-A:10-A:5l0qC:IgG1 | 45 | 17 | 62 | 0.314 | 0.244 | 8C7, ADAM10 binding antibody |
| 13-A:9-A:3cmoH:IgD | 31 | 9 | 40 | 0.280 | 0.255 | YZ18, HIV neutralizing antibody |
| 15-C:9-A:4kvcH:IgD | 55 | 27 | 82 | 0.263 | 0.235 | 2H2, dengue virus binding Fab |
| 15-C:9-A:3hzyB:IgA2 | 39 | 13 | 52 | 0.196 | 0.194 | Chlamydial liposaccharide antigens binding Fab |
| 13-A:12-A:5drwA:IgM | 61 | 81 | 142 | -0.685 | 0.533 | BCR fragment involved in CLL |
| 13-A:10-A:5i66A:IgM | 58 | 78 | 136 | -0.775 | 0.367 | Fab 8B6, monoclonal antibody against serotonin transporter |

**Supplementary Table S3: Characteristics of stable class-switched SCAGs features selected by cross-validated LASSO model for the prediction of food sensitization status.** Sensit and Non-Sensit signify the number of food sensitized subjects and subjects without a food sensitization expressing sequences belonging to any of the 9 class-switched only SCAGs features. Total represents the total number of subjects in our study cohort expressing sequences belonging to each of the 9 class-switched SCAGs features. For each stable class-switch SCAGs feature, their mean weights and standard deviation across folds were calculated. A description of the antibody source of the CDR3 structure is also shown.

| **Class-switched SCAGs** | **Sensit** | **Non-Sensit** | **Total** | **Coefficient mean** | **Std. dev of weights** | **CDR3 Source Description** |
| --- | --- | --- | --- | --- | --- | --- |
| 15-C:9-A:4nzrH:IgG2 | 41 | 10 | 51 | 0.370043302 | 0.251039974 | PGT135, HIV binding antibody (structure not bound to antibody) |
| 13-A:9-A:5xj4H:IgA1 | 50 | 23 | 73 | 0.292572117 | 0.195375965 | Complex structure of durvalumab-scFv/PD-L1 |
| 13-A:10-A:5ds8A:IgG3 | 36 | 11 | 47 | 0.27281385 | 0.196424741 | Hpu98, anti-hypusine antibody |
| 13-A:10-A:5l0qC:IgG1 | 45 | 17 | 62 | 0.266595559 | 0.210753732 | 8C7, ADAM10 binding antibody |
| 13-A:10-A:2op4H:IgG2 | 27 | 5 | 32 | 0.179773069 | 0.178470611 | IG9, quorum-quenching antibody |
| 15-A:9-A:4qwwD:IgE | 11 | 45 | 56 | -0.383559019 | 0.23871017 | Fab410, snake venom binding antibody |
| 13-A:10-A:5mheH:IgA1 | 64 | 82 | 146 | -0.544045723 | 0.43688061 | Anti-T4 antibody |

**Supplementary Table S4: Characteristics of stable VJ3 with isotype features selected by cross-validated LASSO model for the prediction of food sensitization**. Sensit and Non-Sensit signify the number of food sensitized subjects and subjects without a food sensitization expressing sequences belonging to any of the 9 VJ3 with isotype features. Total represents the total number of subjects in our study cohort expressing sequences belonging to each of the 9 VJ3 with isotype features. For each stable VJ3 with isotype feature, their mean weights and standard deviation across folds was calculated.

| VJ3 with Isotype Features | Sensit | Non-Sensit | Total | Weights mean | Std. dev of weights |
| --- | --- | --- | --- | --- | --- |
| 28027:V4-39:J6:IgD | 17 | 2 | 19 | 1.223 | 0.275 |
| 15474:V4-59:J6:IgD | 21 | 3 | 24 | 0.967 | 0.374 |
| 20163:V4-39:J6:IgD | 14 | 1 | 15 | 0.880 | 0.3487 |
| 39077:V5-51:J6:IgD | 14 | 1 | 15 | 0.667 | 0.420 |
| 62796:V5-51:J4:IgD | 13 | 1 | 14 | 0.537 | 0.335 |
| 6220:V4-59:J4:IgD | 11 | 0 | 11 | 0.402 | 0.339 |
| 18:V4-61:J4:IgE | 6 | 28 | 34 | -0.242 | 0.239 |
| 25:V2-26:J4:IgE | 11 | 48 | 59 | -0.893 | 0.304 |
| 139:V4-59:J6:IgE | 5 | 34 | 39 | -0.9256 | 0.272 |

**Supplementary Table S5: Characteristics of stable VJ3 with isotype features selected by cross-validated LASSO model for the prediction of HIV infection status**. HIV-Pos and HIV-Neg signify the number of infected and uninfected subjects expressing sequences belonging to any of the 24 VJ3 with isotype features. Total represents the total number of subjects in our study cohort expressing sequences belonging to each of the 24 VJ3 with isotype features. For each stable VJ3 with isotype feature, their mean weights and standard deviation across folds was calculated.

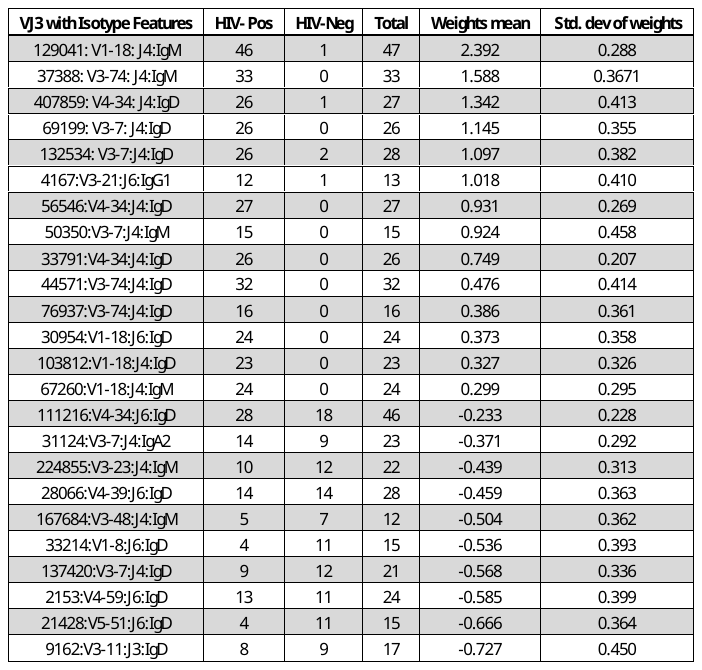

**Supplementary Table S6: Demographic, IgH sequencing statistics of the subjects in HIV cohort.** *Numbers of rearranged IgH sequence reads and number of B cell clones (see Methods), also shown by isotype.

|  |  | HIV infection status | | |
| --- | --- | --- | --- | --- |
|  |  | HIV-Pos | HIV-Neg | all |
| n |  | 101 | 43 | 144 |
| age (years) | minimum | 19 | 20 | 19 |
|  | mean | 32.5 | 29.0 | 31.5 |
|  | maximum | 64 | 51 | 65 |
| sex | female | 66 | 22 | 88 |
|  | male | 35 | 21 | 56 |
| IgH reads* | mean | 266,301 | 106,933 | 218,712 |
| IgM | mean | 42,708 | 17,700 | 35,240 |
| IgD | mean | 59,249 | 29,113 | 50,249 |
| IgA | mean | 47,943 | 18,952 | 39,286 |
| IgG | mean | 54,082 | 21,046 | 44,217 |
| IgE | mean | 62,942 | 21,104 | 50,776 |
| IgH clones* | mean | 75,673 | 41,915 | 65,593 |
| IgM | mean | 25,807 | 12,928 | 21,961 |
| IgD | mean | 34,456 | 19,806 | 30,082 |
| IgA | mean | 7,298 | 5,034 | 6,622 |
| IgG | mean | 11,421 | 5,068 | 9,524 |
| IgE | mean | 176 | 143 | 166 |

**Supplementary Table S7: Demographic, IgH sequencing statistics of the subjects the food sensitization cohort.** *Numbers of rearranged IgH sequence reads and number of B cell clones (see Methods), also shown by isotype.

|  |  | food sensitization status | | |
| --- | --- | --- | --- | --- |
|  |  | non-sensit. | sensit. | all |
| n |  | 72 | 75 | 147 |
| age (years) | minimum | 0.53 | 0.39 | 0.39 |
|  | mean | 3.22 | 2.96 | 3.09 |
|  | maximum | 5.84 | 5.79 | 5.84 |
| sex | female | 37 | 28 | 65 |
|  | male | 35 | 47 | 82 |
| race | black | 56 | 43 | 99 |
|  | non-black | 16 | 32 | 48 |
| IgH reads* | mean | 133,879 | 302,067 | 219,244 |
| IgM | mean | 49,098 | 80,866 | 65,166 |
| IgD | mean | 34,939 | 81,873 | 58,761 |
| IgA | mean | 25,816 | 52,027 | 39,119 |
| IgG | mean | 21,191 | 54,343 | 38,018 |
| IgE | mean | 3,463 | 34,996 | 19,369 |
| IgH clones* | mean | 60,446 | 122,982 | 92,187 |
| IgM | mean | 29,289 | 52,349 | 40,952 |
| IgD | mean | 21,331 | 51,031 | 36,405 |
| IgA | mean | 6,177 | 11,605 | 8,932 |
| IgG | mean | 5,575 | 12,764 | 9,334 |
| IgE | mean | 46 | 286 | 168 |

**Supplementary Figures**

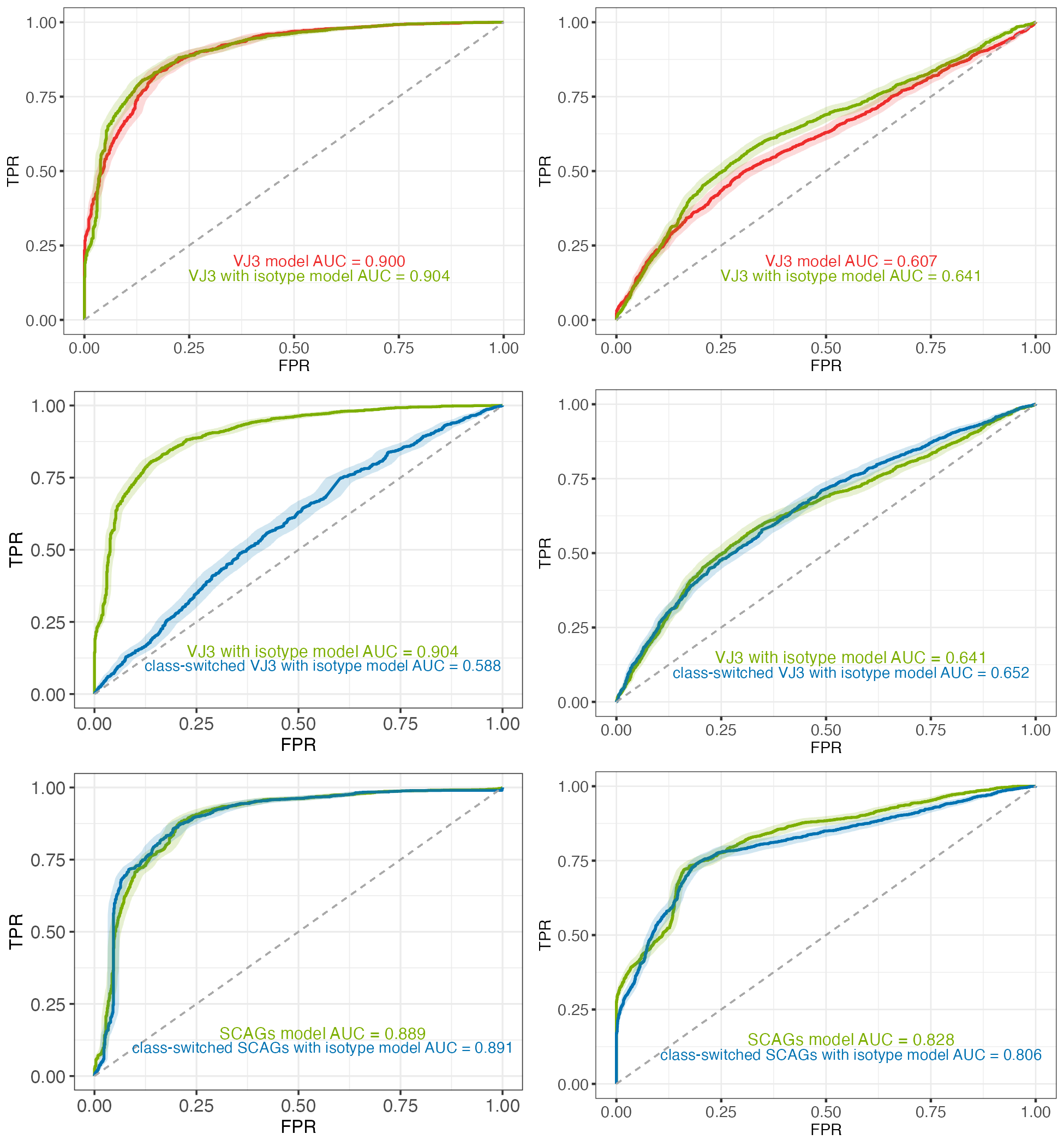

**E**

**B**

**F**

**C**

**D**

**A**

**Supplementary Figure 1: ROC curve and corresponding AUC values comparing VJ3, VJ3 with isotype and SCAGs feature models to their class-switched isotype feature models(A)** HIV status prediction (HIV infected vs. HIV uninfected) using VJ3 features (red curve) and VJ3 with isotype features (green curve). **(B)** HIV status prediction (HIV infected vs. HIV uninfected) using VJ3 with class-switched isotype features (blue curve) and using VJ3 with isotype features (green curve). **(C)** HIV status prediction (HIV infected vs. HIV uninfected) using class-switched SCAGs features (blue curve) and using all SCAGs features (green curve). **(B)** Food sensitization status predictions (food sensitized vs. non-food sensitized) using VJ3 features (red curve) and using VJ3 with isotype features (green curve) **(E)** Food sensitization status predictions (food sensitized vs. non-food sensitized) using VJ3 with class-switched isotype features (blue curve) and using VJ3 with isotype features (green curve). **(F)** Food sensitization status predictions (food sensitized vs. non-food sensitized) using class-switched SCAGs features (blue curve) and using all SCAGs features (green curve). AUC scores highlighting the predictive power of both models ran for both cohorts are shown in blue (class-switched) and light green (VJ3 with isotype or SCAGs). Shaded error regions represent the standard error across all folds. Sensitization status numbers for participants analyzed in the food sensitization cohort were: 70 sensitized and 83 non-sensitized. HIV status for participants in the HIV cohort was: 101 HIV infected and 43 HIV not infected. The axes are the True Positive Rate (TPR) and False Positive Rate (FPR).

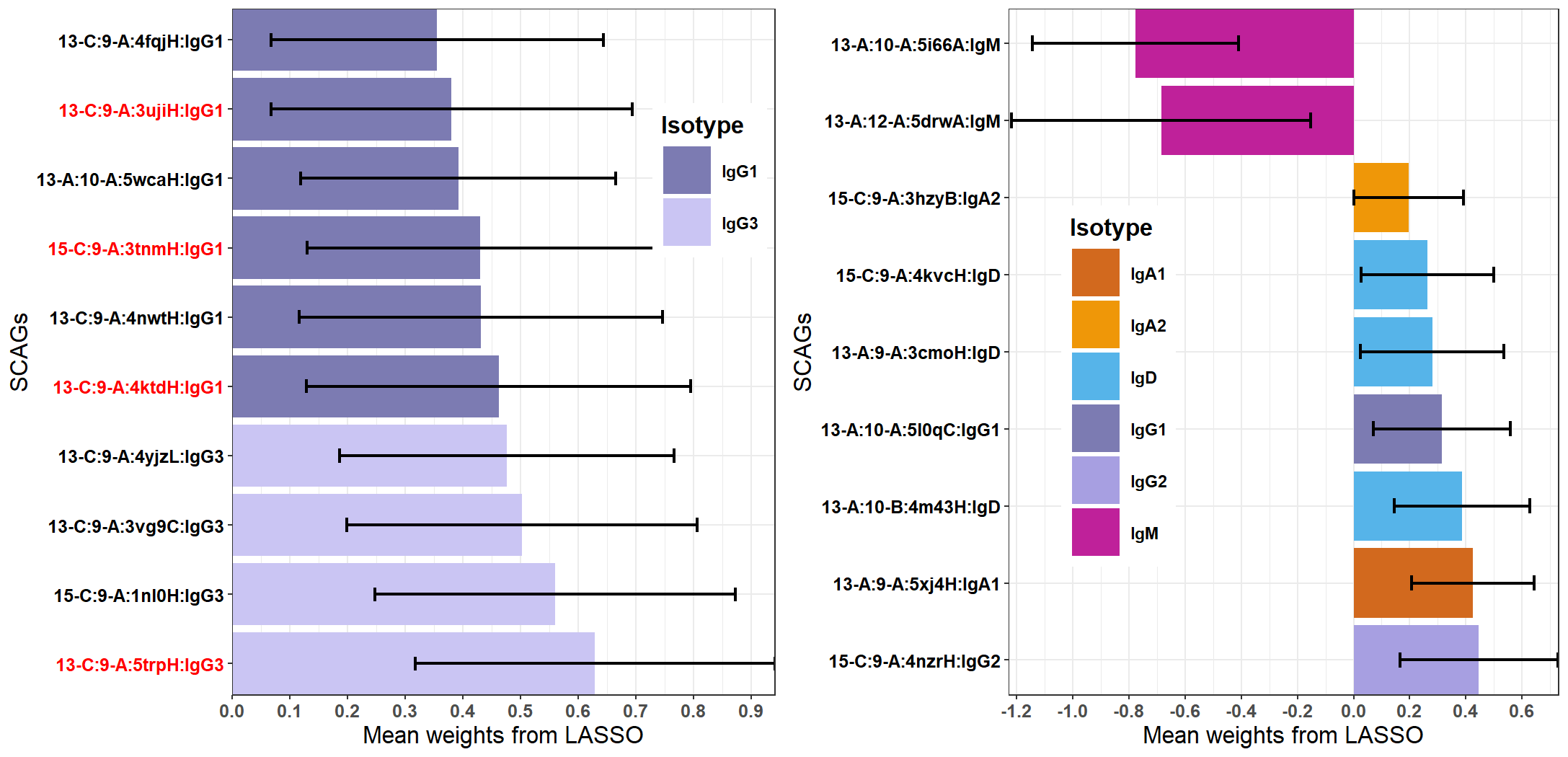

**Supplementary Figure 2: Regression weights for SCAGs features selected by LASSO models.** Weights from four-fold cross-validated LASSO food sensitization status prediction model (food sensitized vs. non-food sensitized) using SCAGs features. Features are labeled based on their assigned CDR1/2 canonical classes, CDR3 PDB chain, and expressed isotype. Error bars represent the range of weights (mean +/- standard deviation) across folds.

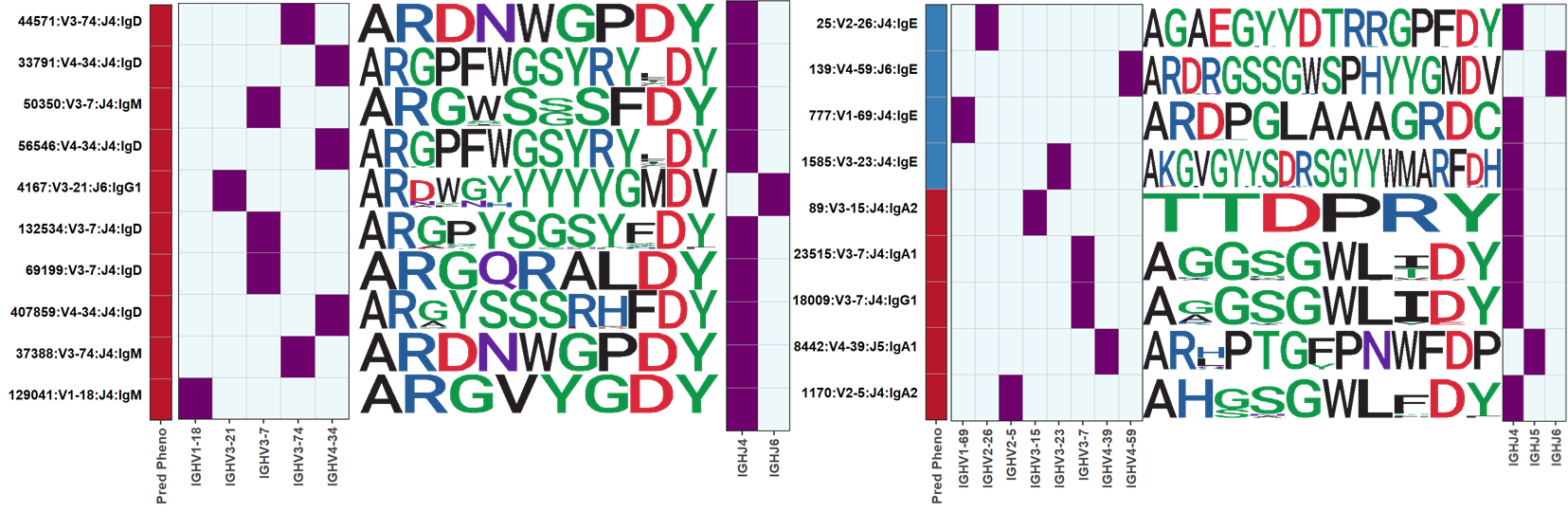
**Supplementary Figure 3: Heatmap plot highlighting V, CDR3 sequence logo and J segments used by sequences in each VJ3 with isotype feature group. (A)** Ten positively associated HIV infection VJ3 with isotype features out of twenty-four. **(B)** Food sensitization class-switched only VJ3 with isotype features. For each feature, the V and J segments usage is represented by a cell in heatmap with higher color intensity indicating higher use of those gene segments by sequence clones which make up the feature. Sequence logo plots represent aligned CDR3 amino acids sequences for the most frequent clones of each feature offering a richer representation of the CDR3’s. The Pred Pheno column represents the association of features with HIV infection or food sensitization (Red: associated with HIV infection or food sensitization, Blue: associated with absence of HIV infection or food sensitization).

**B**

**A**

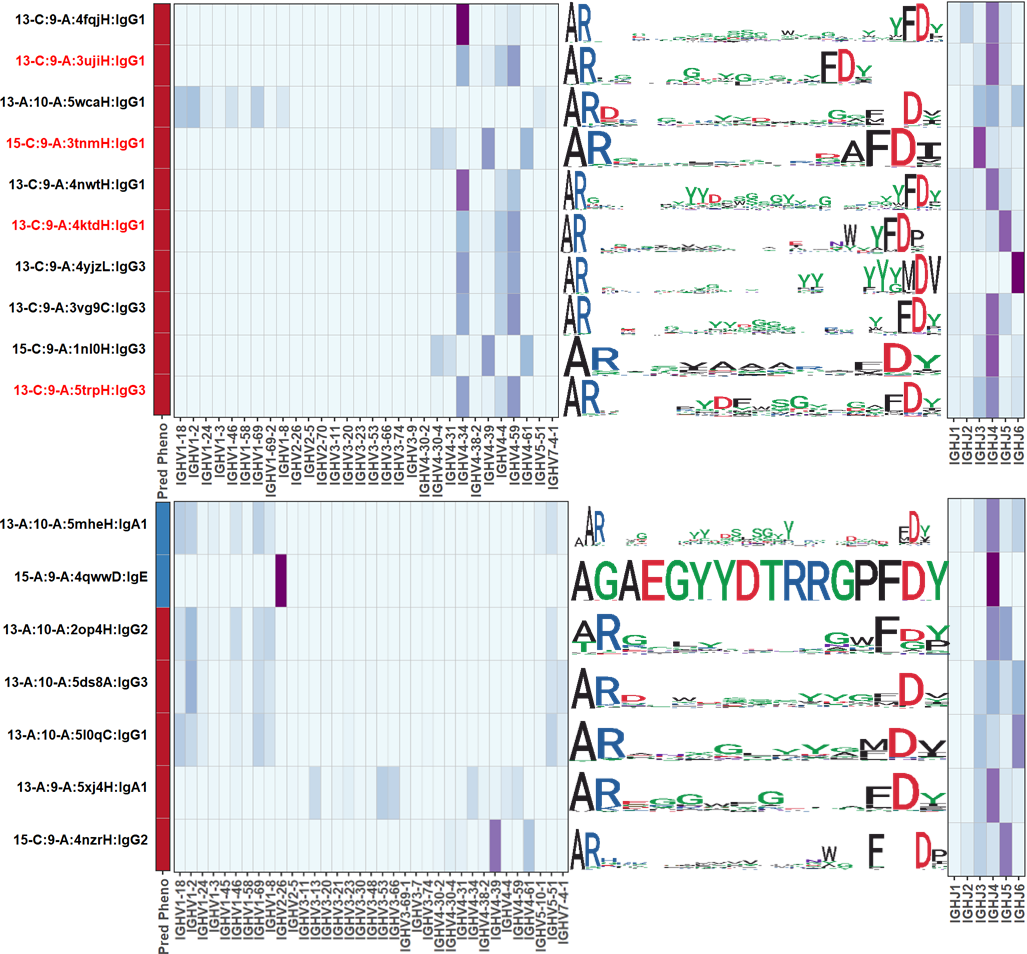

**B**

**A**

**Supplementary Figure 4: Heatmap plot visualizing V, CDR3 sequence logo and J segments used by sequences in each SCAGs feature group. (A)** Ten positively associated HIV infection SCAG features out of twenty-one. **(B)** Food sensitization class switched only SCAG features. For each feature, the V and J segments usage is represented by a cell in heatmap with higher color intensity indicating higher use of those gene segments by sequence clones making up the feature. Sequence logo plots represent aligned CDR3 amino acids sequences for the most frequent clones of each feature offering a richer representation of the CDR3’s. The Pred Pheno column represents the association of features with HIV infection or food sensitization (Red: associated with HIV infection or food sensitization, Blue: associated with absence of HIV infection or food sensitization).

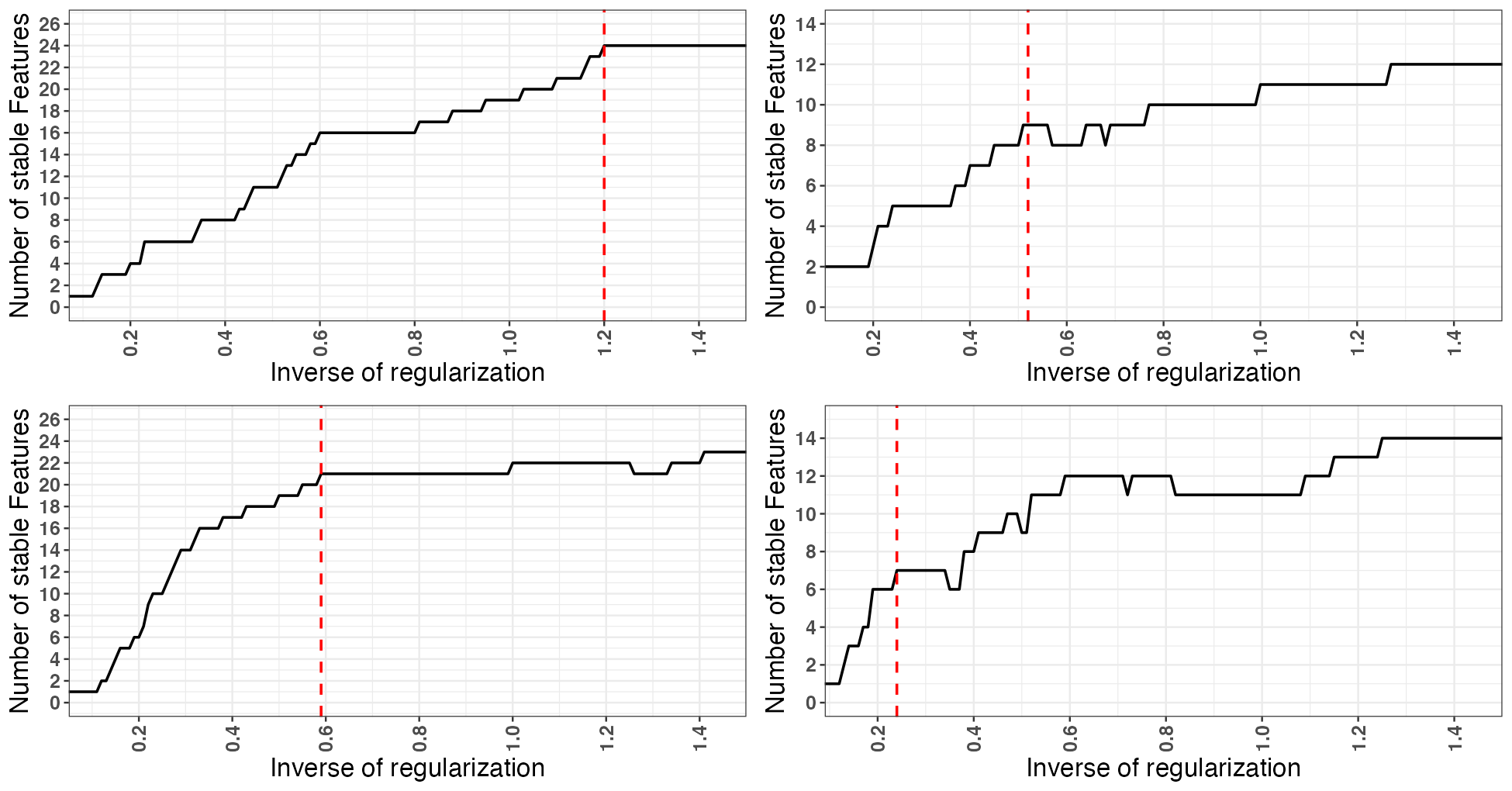

**C**

**D**

**B**

**A**

**Supplementary Figure 5: Representative LASSO model and predictive feature selection for VJ3 and SCAGs model exploration.** Repeated four-fold cross-validated LASSO regression models with varying regularization strengths using VJ3 with isotype, VJ3 with class-switched isotype, SCAGs and class-switched SCAGs feature groups. At each regularization strength, only stable features, i.e. whose mean weight plus-or-minus one standard deviation across all folds did not intersect zero are selected as being consistently included in models across multiple folds. The inverse of regularization strength (x-axis) and its corresponding number of stable features (y-axis) are plotted for **(A)**, HIV infection VJ3 with isotype features **(B**), food sensitization VJ3 with class-switched isotype features. **(C)**, HIV infection SCAG features **(D)**, food sensitization class-switched SCAG features. A final representative LASSO model is selected based on the appearance of local maxima. as indicated by the red dotted lines.
